# Identification of Sample Processing Errors in Microbiome Studies Using Host Genetic Profiles

**DOI:** 10.1101/2025.09.07.674724

**Authors:** Julia Urban, Aya Brown Kav, William F. Kindschuh, Heekuk Park, Raiyan R. Khan, Emily Watters, Itsik Pe’er, Anne-Catrin Uhlemann, Tal Korem

## Abstract

In microbiome studies, sample processing errors are frequent and difficult to detect, especially in large studies involving multiple sites, personnel, and sample types. We present two complementary approaches to identify such errors using host DNA profiled via metagenomic sequencing of microbiome samples. The first approach compares host SNPs inferred from metagenomics to independently obtained genotypes (e.g., microarray genotypes) to match samples to their donors, while the second method compares metagenomics-inferred SNPs between samples to identify samples supplied by the same donor. Furthermore, we demonstrate that combining these methods with experimental metadata provides greater confidence in the identification of errors. Analyzing a longitudinal vaginal microbiome dataset, we demonstrate the ability of our approach to identify mislabeled samples. Using subsampling, we further show that our methods are robust to low sequencing coverage. Overall, our analysis highlights the frequency of processing errors in microbiome studies. We therefore recommend applying error-detection methods in all studies with suitable data.

## Introduction

Modern microbiome studies are large and complex, often spanning hundreds of samples of multiple types and from separate sources. Data generation from microbial samples entails a series of processes from sample collection to library preparation, with each potentially introducing errors such as sample mislabeling or swapping. Each stage may involve different personnel or even separate locations in the case of multi-site studies, amplifying the risk of error. While these errors are likely to be common, they are difficult to detect and can obscure associations if left uncorrected.

Microbiome samples typically contain substantial fractions of host DNA^1^. Studies that profile the human microbiome via shotgun metagenomics could potentially leverage sequencing reads originating from the human host, ranging from 5-10% in fecal samples to >85% in vaginal samples^1^, to identify and even correct errors long after sample processing has completed. This could be achieved using independently acquired host genotypes — for example, in a study of the mouse gut microbiome, comparison of metagenomics to host microarray profiling enabled successful matching of samples to their donors^2^. Similar approaches have been used to compare different human sequencing modalities, as well as the same modality across samples^3–6^. While most microbiome studies do not include host genotyping or sequencing, many include multiple samples from the same host. In such cases, genotypes can be inferred from metagenomic sequencing and compared to confirm similarity between samples from the same donor.

Here, we developed methods for both approaches and confirmed their ability to detect sample processing errors from metagenomic sequencing data, even at low coverage of the human genome.

Furthermore, we showed that combining the results of our methods with experimental metadata (e.g., co-localization of samples in processing batches) can increase confidence in their conclusions and potentially also reveal the source of errors. Analyzing a longitudinal vaginal metagenomic cohort with paired microarray genotyping, we demonstrated that sample processing errors are common and correctable. We therefore recommend that microbiome studies implement such processing-error detection methods as standard quality control procedures.

## Results

### Host genotype-based processing-error detection

To detect sample processing errors in microbiome studies, we developed two complementary methods which rely on calling host Single Nucleotide Polymorphisms (SNPs) from metagenomic sequencing.

One method makes “metagnome:genotype” comparisons, i.e., compares the genotypes inferred from metagenomics to genotypes from an independent source (e.g., microarrays). The second method makes “metagenome:metagenome” comparisons, comparing genotypes inferred from two metagenomic samples to identify samples likely supplied by the same host (**Methods**; **Fig. 1**). To ensure SNP quality and reduce the effects of sequencing error and ambiguous read mapping, we excluded repeat regions of the human genome and retained only sequenced bases with high sequencing or alignment quality scores (**Methods**). We developed a SNP concurrence score and thresholds that could then be used to identify samples likely supplied by the same individual (**Fig. 1**).

**Figure 1.**
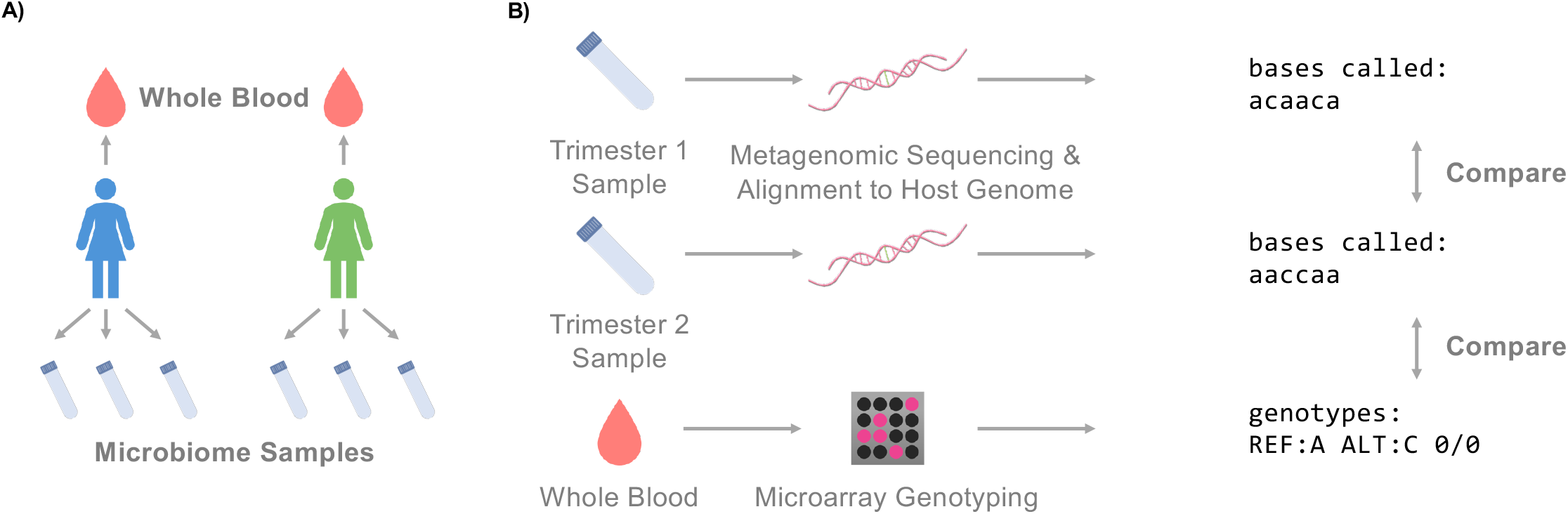
Detection of sample processing errors using metagenome-inferred host SNPs. **(A)** Each participant in the nuMoM2b study supplied whole blood for genotyping in addition to up to three vaginal swabs. **(B)** Human SNPs inferred from metagenomic sequencing could be compared between vaginal samples from the same individual to confirm that the samples indeed came from the same individual. SNPs inferred from metagenomics can also be compared to SNPs obtained via microarray genotyping to match samples to their likely donors.

To demonstrate these methods, we applied them to four batches of vaginal metagenomic samples (**Methods**), which included 250 samples from 108 participants, which were a subset of a larger cohort of 10,038 individuals^7^. Nine, 56, and 43 individuals supplied one, two, and three samples, respectively, with 99 collected during each of the first two research visits (6-14 and 16-22 weeks of gestation), and 52 during the third visit (22-30 weeks of gestation). Three samples failed sequencing (less than 200K read pairs before processing) and were thus excluded (**Fig. S1**). Samples were sequenced to a mean±std depth of 36.9 ± 13.2 million read pairs, with 34.4 ± 12.3 million read pairs mapping to the human genome (GRCh38), corresponding to 2.12x ± 0.75x coverage. Host genotypes, measured via microarrays, were available for 9,756 individuals from the larger cohort, including 106 of 108 participants whose vaginal metagenomics were analyzed here (**Methods**).

### Comparison of metagenomics-derived and independently obtained genotypes identifies processing errors

To identify the donor of each of our 247 vaginal samples, we inferred genotypes from metagenomic sequencing and compared them to all available array-derived genotypes (9,756 individuals). We quantified the similarity between SNP profiles in each metagenome:genotype comparison using a concurrence score, calculated as the percentage of shared SNPs for which metagenomics-inferred alleles were a subset of those derived from independent genotyping (“lax concurrence”; **Methods**). As expected, comparison between metagenomes and genotypes labeled for the same donor (“matched” comparisons, **Fig. S2A**) yielded substantially higher concurrence scores than comparisons labeled for different donors (“mismatched” comparisons; **Fig. 2A**). We selected an operational threshold to identify sample processing error, 98%, which was the midpoint between the upper and lower outer fence of the mismatched and matched distribution, respectively (**Figs. 2A, S2A**). In three of the four sample batches (batches 2-4), only matched comparisons yielded concurrence scores above 98%, and only mismatched comparisons yielded scores below this threshold, indicating that samples had been labeled and processed correctly (**Fig. 2B**).

**Figure 2.**
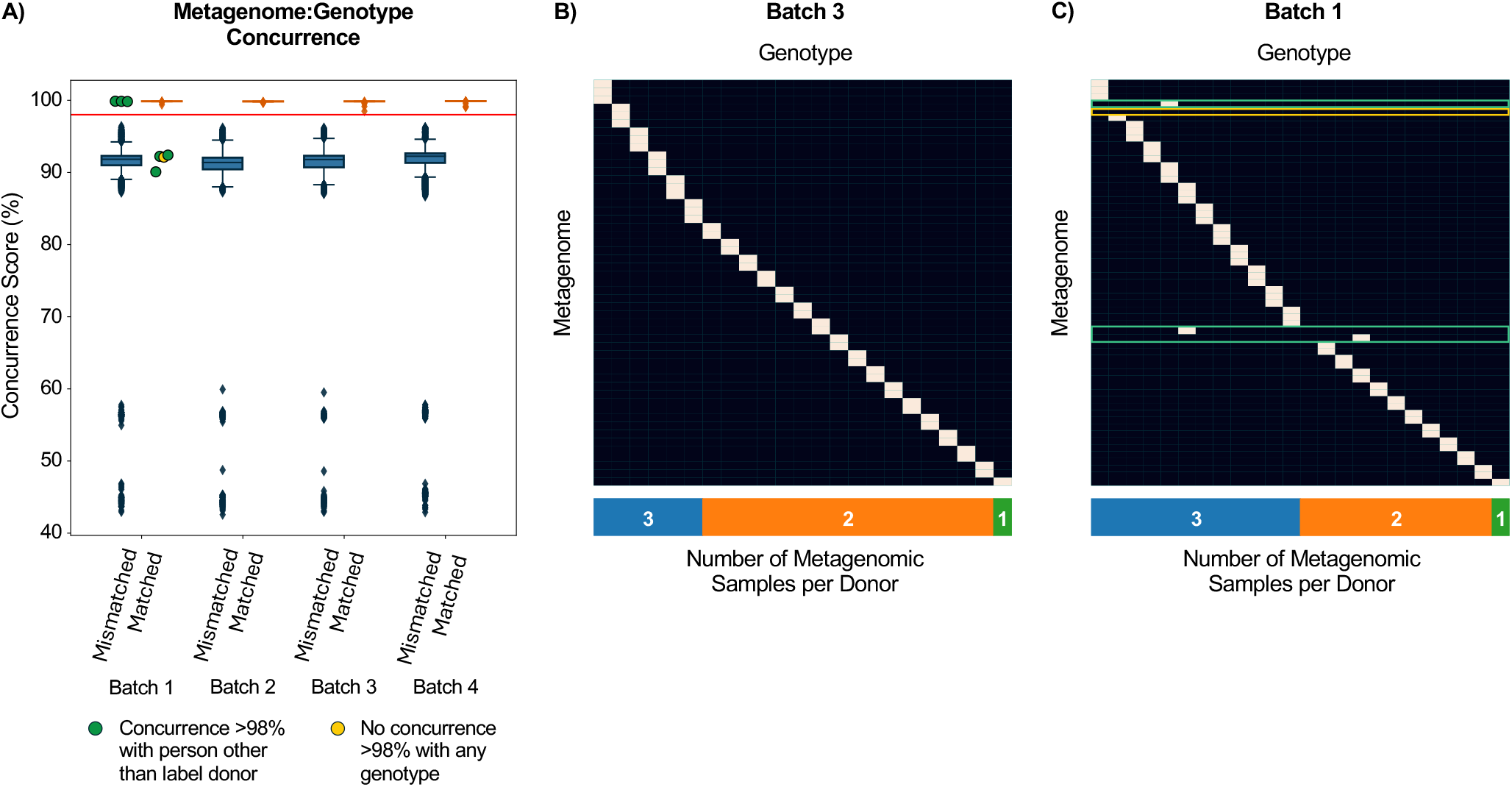
Concurrence score differentiates matched from mismatched metagenome:genotype pairs. **(A)** Boxplots (Box, IQR; line, median; whiskers, 1.5xIQR) showing concurrence score for all 247 metagenomes compared to all 9,756 available microarray genotypes. Comparisons of metagenomes to the microarray genotypes of their labelled donor (“matched”) are plotted separately from metagenomes compared to the microarray genotypes of different individuals (“mismatched”). Our operational 98% concurrence score threshold is marked in red. Green dots mark samples with concurrence >98% with individuals other than their labeled donors; the yellow dot marks a sample that did not have high concurrence with any microarray genotype. **(B-C)** Heatmaps showing metagenome:genotype comparisons above (white) and below (black) the 98% threshold for batches 3 (B) and 1 (C). Both heatmaps were subset to individuals with at least one sample in the relevant batch. “Off-diagonal” high concurrence scores indicate that a sample’s donor is an individual other than the labelled donor (green outline), while a row with no comparisons above 98% concurrence indicates that a metagenome did not have high similarity to the genotypes of any individual with samples in the same batch (yellow outline).

In batch 1, four metagenomic samples had concurrence scores below 98% (range of 90.08-92.40%) in matched comparisons with the genotypes of their labeled donors. Of these, three had concurrence scores above 98% with the genotypes of other individuals (“mismatched comparisons”; 99.87, 99.85, and 99.83%; green points in **Fig. 2A**; green outlines in **Fig. 2C**). All three samples were thus implicated in a possible sample mislabeling. The fourth sample did not resemble any of the 9,756 nuMoM2b genotypes (yellow point in **Fig. 2A**; yellow outline in **Fig. 2C**), potentially because it originated from one of the two individuals in our dataset who were not genotyped.

### Comparison of metagenomics-derived SNPs from the same host reveals processing errors

In microbiome datasets lacking independent genotyping (e.g., datasets without microarray genotypes), metagenome-inferred SNPs can be compared between multiple samples from the same host to confirm that they are similar to one another. Even within our four batches, two individuals with longitudinal samples lacked microarray data, exemplifying the utility of such an approach. We therefore compared metagenome:metagenome sample pairs, and calculated a slightly different concurrence score - the percentage of shared SNPs in which the inferred alleles from both samples match (“strict concurrence”; **Methods**). Once again, metagenome:metagenome comparisons from the same labeled donor (“matched” pairs; **Fig. S2B**) had substantially higher concurrence scores than pairs from different donors (“mismatched” pairs; **Fig. 3A**). Here we selected a different operational threshold, 90%, which was again the midpoint between the lower and upper outer fence of the matched and mismatched distributions, respectively (**Figs. 3A, S2B**). As for metagenome:genotype comparisons, our metagenome:metagenome comparisons gave no indication of processing errors in three of four batches, as the metagenome of each sample resembled only samples from the same individual (**Fig. 3B**).

**Figure 3.**
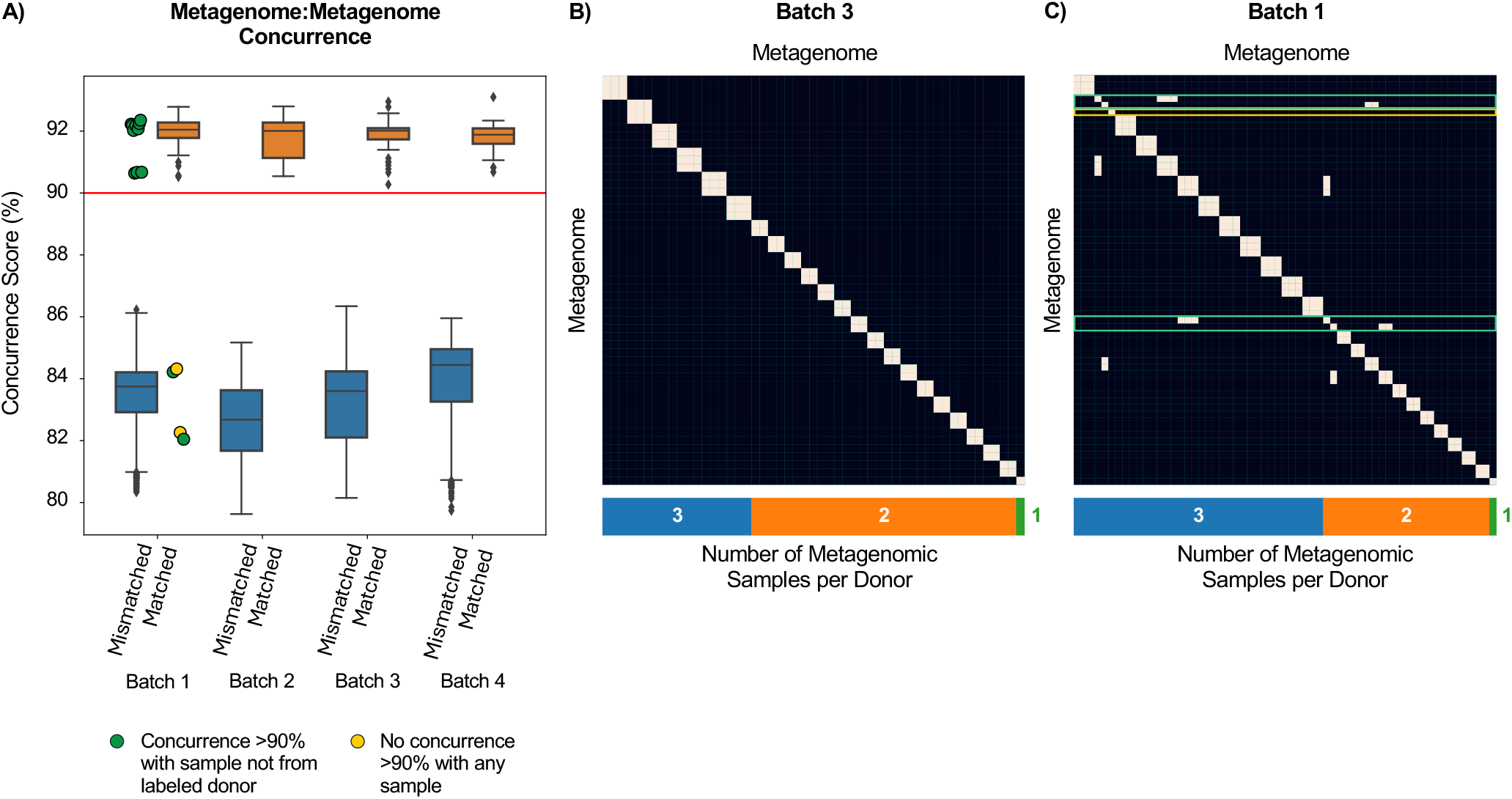
Concurrence score differentiates metagenomes from different donors. **(A)** Boxplots (Box, IQR; line, median; whiskers, 1.5xIQR) showing concurrence scores for comparisons between metagenomes from the same donor individual (“matched”) versus metagenomes from different individuals (“mismatched”). Our operational 90% concurrence score threshold is marked in red. Green points mark samples with low concurrence to samples from their labeled donor, but high concurrence with samples from other individuals. The two yellow points mark matched comparisons between a sample that had no concurrence >90% with any sample, including two samples labeled as supplied by its donor. **(B-C)** Heatmaps showing metagenome:metagenome comparisons above (white) and below (black) the 90% threshold for batches 3 (B) and 1 (C). The expected case where all samples from the same individual are similar to one another is seen as “boxes” along the diagonal. Off-diagonal high concurrence indicates a processing error, such as mislabeling (green outline). Samples with low concurrence to samples from the same donor (by label) and to all other samples are outlined in yellow.

Similarly, in a metagenome:metagenome analysis of batch 1, the same samples that did not resemble their donors’ microarray genotypes in metagenome:genotype comparisons (**Fig. 2**) also did not resemble other samples from the same donor. Specifically, metagenome:metagenome comparisons identified four samples with concurrence of <90% to other samples from their labeled donor (range of 82.04-84.32%), but >90% to samples from a different individual (range of 90.64-92.35%), providing further evidence of sample processing errors (green points in **Fig. 3A**; green outlines in **Fig. 3C**). A fifth sample in batch 1 did not resemble any other samples, including the two other samples supplied by its labeled donor (yellow points in **Fig. 3A**; yellow outline in **Fig. 3B**). However, these two samples were part of the four samples with high concurrence to samples from a different individual in metagenome:metagenome comparisons and were thus likely mislabeled (topmost green outline in **Fig. 3C**).

### Concurrence analysis is robust to low sequencing coverage

We next evaluated the coverage necessary for robust concurrence analysis. We subsampled alignments to the human genome of all 240 samples with coverage ≥1x to depths ranging from 1K to 2M alignments, simulating 6.2×10^−5^ - 0.12x coverage of the human genome, which we repeated ten times per alignment depth. We then performed concurrence analysis and evaluated whether it indicated similar findings as the original, full-depth analysis, i.e., identical classifications above and below the operational concurrence thresholds.

Even at 75K read pairs (0.0047x coverage), metagenome:genotype comparisons were still accurate, with 100% accuracy for all 10 replicates (**Figs. 4A, S3A-C**). Accuracy began to decrease at depths below 75K read pairs; however, analysis at 50, 25, and 10K read pairs made an average±std of only 0.2±0.42, 2.0±1.76, and 55.0±16.01 mistakes out of over 2.3 million total comparisons, respectively. Metagenome:metagenome comparisons were more sensitive to coverage: though our analysis distinguished metagenome pairs from the same donor from pairs from separate donors at 0.11x coverage (1.75M read pairs) with 100% accuracy across all replicates, accuracy decreased at lower coverage (**Figs. 4B, S3D-F**). Accuracy declined steeply between 0.5M and 0.25M read pairs (0.031x and 0.015x coverage, respectively), falling from 95.06 ± 0.15 to 80.71 ± 0.43%. At 0.1M read pairs or about 0.0062x coverage, only 60.27 ± 0.26% of comparisons yielded the same results as the original, full-depth analyses.

**Figure 4.**
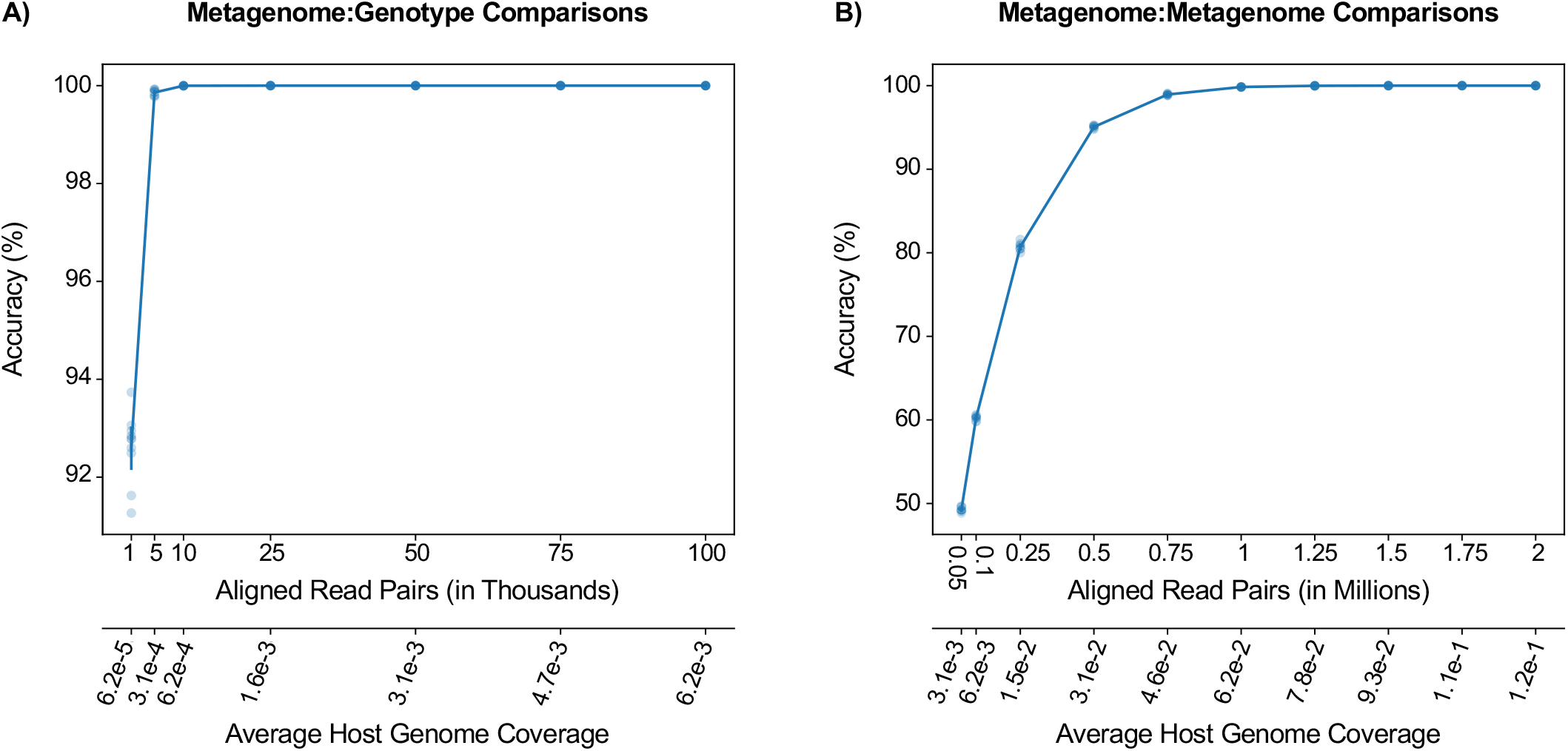
Concurrence analysis is robust for low sequencing coverage. Results are from ten replicates for each subsampling depth. **(A)** Accuracy of metagenome:genotype comparisons on samples subsampled to 1-100K read-pair alignments to the human genome (top x-axis; shown also in units of human-genome coverage on the bottom x-axis). **(B)** Similar to A, showing the accuracy of metagenome:metagenome comparisons on samples subsampled to 50K-2M read alignments. For A and B: Line, average; bars, standard deviation; points, results from individual random replicates. See also **Fig. S3** for similar figures showing auPR and read alignments in log scale.

### Analyzing sample location during processing facilitates high-confidence identification of the cause of processing errors

The spatial and temporal colocation of samples is related to the occurrence of processing errors, as well as other issues such as well-to-well leakage^8,9^. For example, samples processed or stored together may be more easily swapped with one another than samples in separate groupings. Thus, joint investigation of experimental study design alongside concurrence analysis results can elucidate the causes of mistakes. In our analysis, four potential sample processing errors had occurred in batch 1. We therefore used recorded sample positions in the 96-well plate during DNA extraction to visualize sample identity by label and by concurrence analysis (**Fig. 5**). We found that all four potentially mislabeled samples were positioned in the same column, column 3. Furthermore, the metagenome-inferred SNPs of three of these samples (C3, E3 & F3) had high concurrence with the genotypes of the individuals who donated the samples immediately to their right (lax concurrence scores of 99.83, 99.85, and 99.87%, respectively). This provides a strong indication that samples from the adjacent column (column 4) were pipetted twice, causing a sample duplication (**Fig. 5**). Though we suspect that sample D3 was also duplicated, we could not confirm this with metagenome:genotype comparisons, as the donor of the sample to the right of D3 was not genotyped. To further investigate this last remaining sample, we analyzed metagenome:metagenome comparisons. We found that sample D3 resembled samples G5 and H5 with strict concurrence scores of 92.18 and 92.02%, respectively. As G5 and H5 were supplied by the same donor as D4, these comparisons implicated D3 as a duplicate of the sample in well D4, as part of a four-sample duplication event. Overall, this analysis demonstrates that joining concurrence analysis with records of study designs enables identification of causes of processing errors.

**Figure 5.**
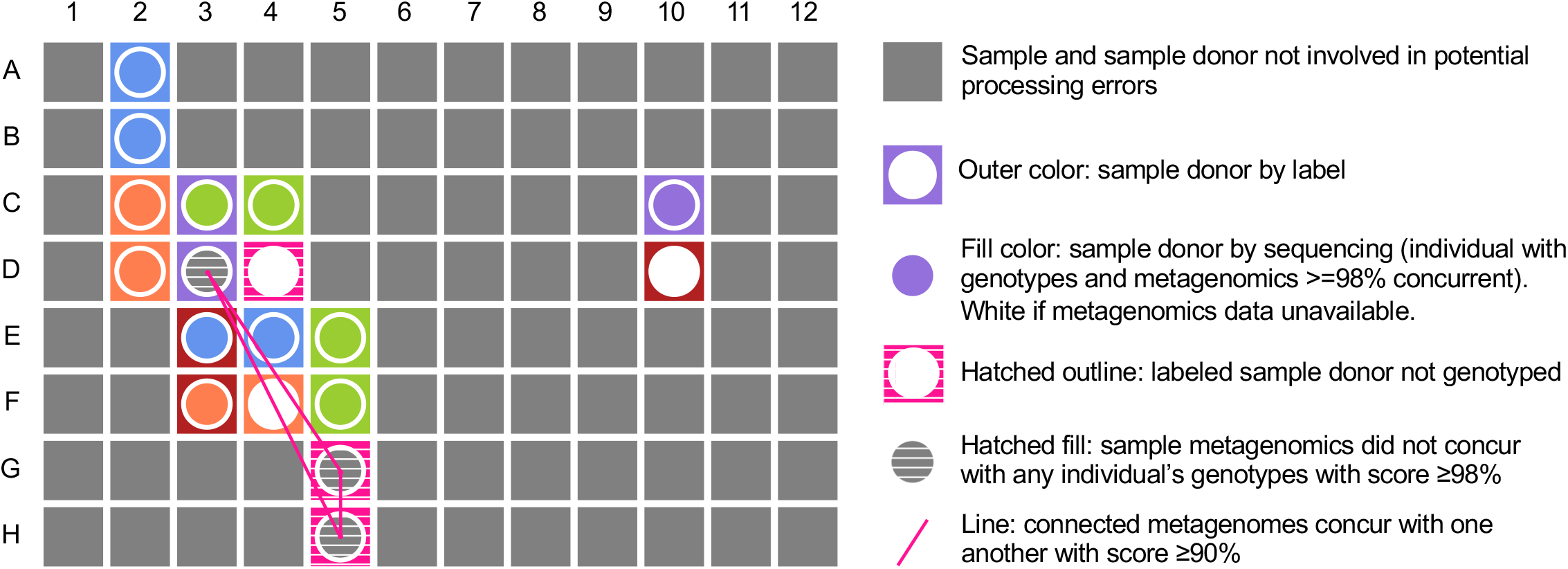
Combining experimental design parameters with concurrence analysis elucidates likely cause for sample processing errors. Shown are recorded positions of samples from batch 1 on a 96-well plate during DNA extraction. The outline color of each well represents the study participant by label and the fill color represents the study participant identified by concurrence analysis with microarray data. Wells in grey represent samples with no involvement in potential processing errors. A possible sample duplication is shown in column 3, rows C-F. For three of these four samples, the sample donor inferred by metagenome:genotype comparisons is actually the donor of the sample directly to the right, rather than the donor listed on the sample’s label. Though sample D3 did not resemble any of the 9,756 available microarray genotypes, the individual who supplied the sample in the well to its right (D4) was not genotyped. Metagenome:metagenome analyses confirmed similarity between D3 and other samples from this individual (G5 and H5; pink lines), thus implicating D3 as the fourth sample involved in the duplication event.

We note that, just as metagenome:genotype comparisons are limited by the availability of independent genotyping data, metagenome:metagenome comparisons are limited by the quantity of available samples per study participant. For example, metagenome:metagenome comparisons alone could not show that sample C10 had been properly labeled, as it did not resemble either of the two samples labeled as belonging to the same donor (C3 and D3). By incorporating metagenome:genotype comparisons, we could confirm that these two samples (C3 and D3) were likely mislabeled, and that C10 has high concurrence with the genotype of its labeled donor. This further demonstrates the complementarity of the different analyses that we performed, and how they benefit from joint analysis with experimental parameters.

## Discussion

Sample processing errors are expected in microbiome studies, especially as they become larger, more complex, and include multi-omic measurements. We demonstrate that some of these errors could be identified by comparing host genotypes inferred from metagenomic sequencing data either to genotypes inferred from other samples from the same host, or to those acquired independently. We show that such comparisons are robust to low sequencing depth, and that inference could be improved even further by considering experimental design parameters. We demonstrate the utility of our approach using a vaginal metagenomic dataset paired with genotypes obtained from microarrays, in which we confidently identify a duplication event affecting four samples. Our results highlight the frequency of sample processing errors in microbiome studies. We advocate for the implementation of error-detection methods such as those described here in any microbiome that has the necessary data.

In some cases, our approach could be limited by coverage, especially in the case of metagenome:metagenome comparisons, which we showed to decrease in accuracy below 1.75M read pairs aligned to the human genomes (or 0.11x coverage). Our results indicate that a vaginal microbiome study, in which over 90% of reads would be human^10^, would require 2M and 84K total read pairs for accurate metagenome:metagenome and metagenome:genotype analyses, respectively. A stool microbiome study, in which approximately 10% of reads would be human, would require approximately 17.5M and 750K total read pairs. These estimates may not be accurate in cases where there is a technical or biological reason for low sequencing depth in a particular sample that is not captured by our subsampling framework.

We note, however, that in many cases, samples with insufficient coverage for concurrence analysis may have such a low sequencing depth that they would in any case be discarded from downstream analyses. Additionally, some metagenomes may fail to reach a concurrence score of 98% with any individual’s genotypes because they are actually a mixture of two or more samples (from well-to-well leakage^8,11^ or accidentally pipetting in the same position twice). In this case, the samples should be excluded from the dataset regardless of the cause underlying their lack of concurrence to the expected individuals’ genotypes.

As we demonstrate here, combining concurrence analysis with analysis of experimental parameters may result in high-confidence identification of the types and sources of the errors that occurred. In such cases, processing errors may be retroactively “fixed” by re-labeling samples, improving the quality of the final dataset. For example, in our processing batch 1, we can relabel the duplicated samples to their correct donor, rather than discard their data.

Our analyses highlight the importance of recording detailed, high-quality sample processing metadata, such as plate and storage box layouts, which are also useful for other goals, such as identification and removal of contamination or handling of processing biases^8,9,12^. However, we note that because human reads are typically removed from microbiome datasets prior to uploading to an online repository, the responsibility of error detection lies with a study’s original authors. Further consideration should be given for ethics approvals for analyses of human genetic data, although in the case of studies with concurrent host genetic data, such approvals are likely already in place.

Sample processing errors, including mixtures, duplications, and swaps are likely common, and may obscure signals between the microbiome and various phenotypes and diseases. Following the approaches described herein, microbiome studies with metagenomic sequencing and independently obtained host genotypes, as well as studies with multiple metagenomic samples per study participant, have ample recourse to detect and address such errors.

## Methods

### nuMoM2b cohort

We analyzed metagenomic sequencing data of 250 banked vaginal samples collected as part of nuMoM2b study^7,13^ paired with previously published microarray data^14^. This analysis was approved by Columbia University’s Institutional Review Boards (IRBs), approval number AAAT7071. Participants were enrolled in the first trimester of pregnancy and self-collected vaginal swabs at up to three research visits (6-14, 16-22, and 22-30 weeks of gestation). Each swab was subsequently suspended in 1 mL 10 mM Tris-Hcl, 0.1 mM EDTA. Included in this analysis are samples from four processing batches, totaling 250 samples from 108 individuals.

### Metagenomic sequencing of vaginal swabs

We used the QIAcube HT and the QIAmp 96 Virus QIAcube HT kit (Cat. No. 57731) to extract DNA from 200 µL aliquots. We prepared sequencing libraries using the Nextera DNA Library Prep Kit (Cat. No. 20060059). We performed paired-end sequencing (2×100bp) on NovaSeq 6000 to a mean±std depth of 36.4 ± 13.7 million read pairs per sample.

### Host genotypes

We obtained genotyping data for 9,757 nuMoM2b participants from Khan et al.^14^, which was conducted on blood samples using the Infinium Multi-Ethnic Global Array (Illumina, USA), containing >1 million variants that are on average 1.4 Kb apart. Standard quality control filters were conducted by Khan et al., using GenomeStudio v2.4 (Illumina), and genotype calls for the ~1.7 million loci that passed initial quality control (98.3% of all markers in the array) were made with Beeline autoconvert (Ilumina).

### Metagenomic sequencing preprocessing

We performed standard quality filtering using Trimmomatic^15^ (v0.39), removing reads containing Illumina adapter sequences, bases at the edges of reads with quality scores <25, and reads of length <50 bp (ILLUMINACLIP:<adapter fasta> LEADING:25 TRAILING:25 MINLEN:50). We aligned filtered reads to the human genome (GRCh38, GenBank assembly GCA000001405.15) using Bowtie2 (v2.5.3)^16^ and calculated sequencing coverage for each sample using samtools (v1.17)^17^. To enrich the dataset for informative hypervariable regions, we filtered our metagenomic alignments to the same loci profiled by microarray. We additionally excluded loci in genomic repeat regions in order to remove ambiguously called SNPs^18^. To distinguish sample processing errors from sequencing or alignment errors, we retrained only sequencing base calls with high sequencing and alignment quality (Phred>36 for both).

### Metagenomics-genotype comparisons

To compare metagenomics to genotype data, we calculated the number of loci shared between them, and, among these, the percentage of loci with concurrent genotypes between metagenomics and microarray. Because microarrays provide reliable information on both alleles, our definition of concurrence in this case is “lax”: a metagenome only has to have a subset of the genotyped alleles to be considered “concurrent” with genotyping at a position. For example, if an individual has the genotype A/G called by microarray at a locus, metagenomic sequencing reads could show only A, only G, or a combination of A and G at the locus to be considered concurrent.

### Metagenomics-metagenomics comparisons

Similarly, to compare two metagenomics samples, we followed a similar calculation, except our definition of concurrence was “strict”. Due to the lower reliability of metagenomics-based genotype calls, both samples had to have the same bases read at a locus to be concurrent at that locus. For the previous example of an individual with an A/G genotype called from one sample, the other sample’s metagenomic sequencing reads would need to contain both A and G at this locus (but no other bases) to be considered concurrent.

## Code availability

All Python code required for conducting and visualizing host SNP comparisons is available on Github https://github.com/korem-lab/metagenome_matcher/.

## Acknowledgments

We thank members of the Korem and Uhlemann groups for useful discussions. We thank all participants, staff, and investigators involved in sample collection and generation of data used in this study. This study was supported by grant funding from the Eunice Kennedy Shriver National Institute of Child Health and Human Development (NICHD): R01 HD106017 (T.K.), R01 HD114715 (T.K.), 5F30 HD108886 (W.F.K.), F31 HD115394 (J.A.U.), U10 HD063036, U10 HD063072, U10 HD063047, U10 HD063037, U10 HD063041, U10 HD063020, U10 HD063046, U10 HD063048, and U10 HD063053; the National Library of Medicine: T15 LM007079 (J.A.U.); and the Clinical and Translational Science Institutes: UL1 TR001108 and UL1 TR000153.

## Author contributions

J.A.U and T.K conceived and designed the study. J.A.U. developed methods and analyzed data, with assistance from A.B.K, W.F.K, R.R.K, and I.P. H.P. and E.W. prepared and sequenced the microbial samples under supervision by A.-C.U. A.B.K., J.A.U., and W.F.K. processed sequencing data. J.A.U. and T.K. wrote the manuscript with critical input from all authors. T.K. supervised the study.

## Supplementary Material

**Supplementary Figure 1.**
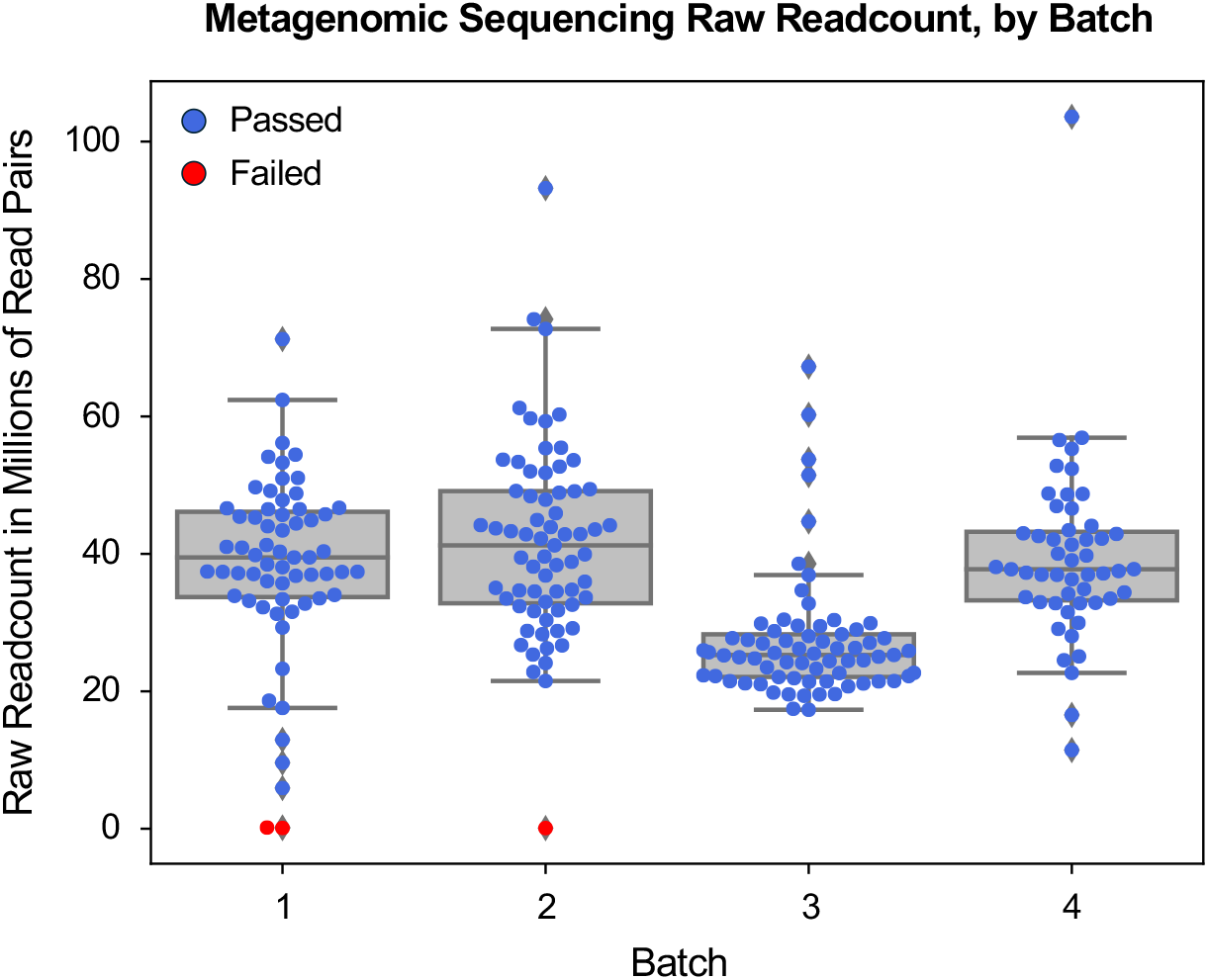
Raw readcount of metagenomic samples, by batch. Three samples failed metagenomic sequencing with raw readcounts < 1M read pairs (91,462, 109,247, and 178,358 read pairs; samples marked with red points) and were thus removed prior to downstream analyses.

**Supplementary Figure 2.**
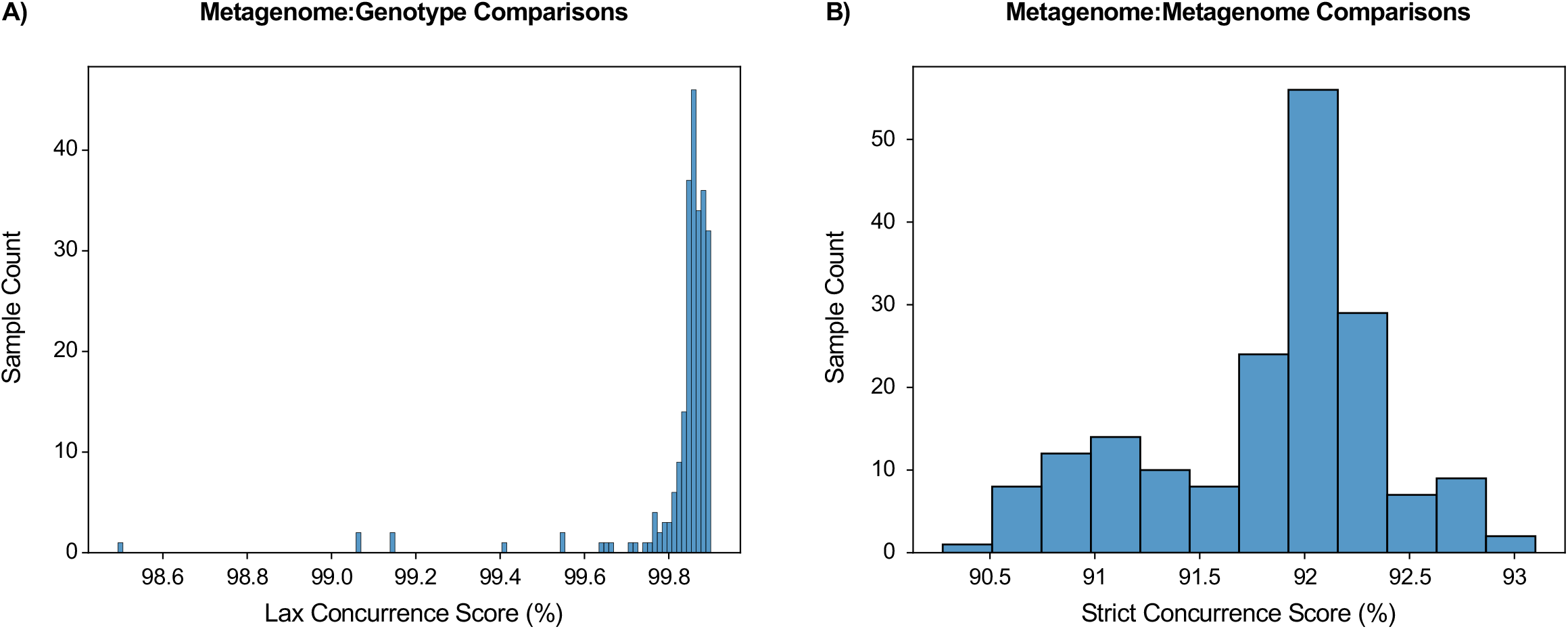
Score distributions for comparisons that are not indicative of sample processing error. **(A)** Lax concurrence score distribution for matched metagenome:genotype comparisons (comparisons of metagenomes with their donors by label) with concurrence >98%, our operational threshold for determining true donor identity. **(B)** Strict concurrence score distribution for metagenome:metagenome comparisons between samples from the same donor with concurrence >90%, our threshold for identifying samples from the same donor.

**Supplementary Figure 3.**
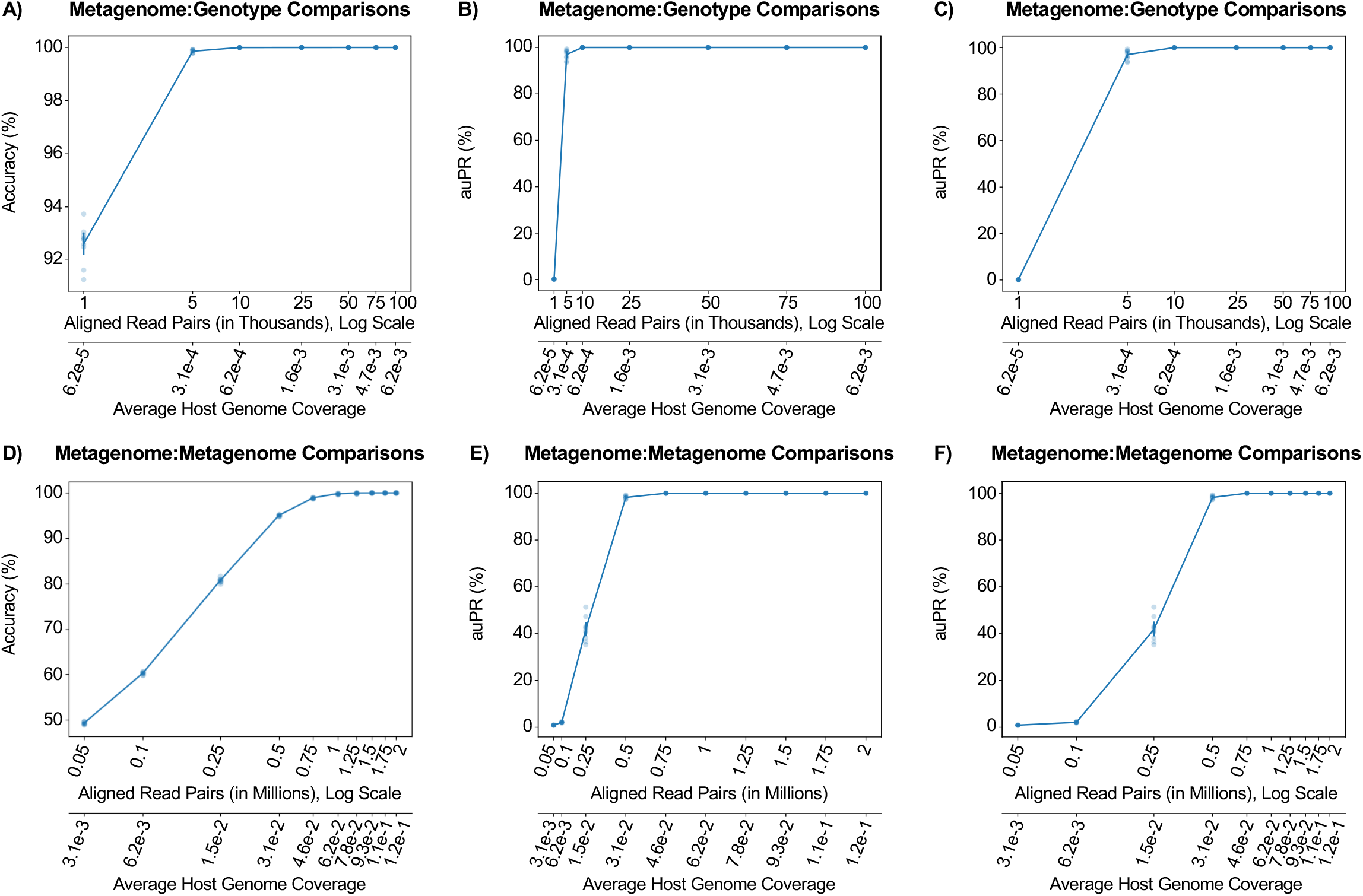
Concurrence analysis is robust for low sequencing coverage. Same as **Fig. 4**, but with read count displayed in log-scale to better show accuracy at lower read thresholds, as well as area under the precision-recall curve (auPR) instead of accuracy.

## References

1. Pereira-Marques, J. et al. Impact of host DNA and sequencing depth on the taxonomic resolution of whole metagenome sequencing for microbiome analysis. Front. Microbiol. 10, 1277 (2019).

2. Lobo, A. K. et al. Identification of sample mix-ups and mixtures in microbiome data in Diversity Outbred mice. G3 (Bethesda) 11, jkab308 (2021).

3. Lee, S. et al. NGSCheckMate: software for validating sample identity in next-generation sequencing studies within and across data types. Nucleic Acids Res. 45, e103 (2017).

4. Chun, H. & Kim, S. BAMixChecker: an automated checkup tool for matched sample pairs in NGS cohort. Bioinformatics 35, 4806–4808 (2019).

5. Eschrich, S. A., Yu, X. & Teer, J. K. Fast all versus all genotype comparison using DNA/RNA sequencing data: method and workflow. BMC Bioinformatics 24, 164 (2023).

6. Chu, J., Rong, J., Feng, X. & Li, H. Ntsm: An alignment-free, ultra-low-coverage, sequencing technology agnostic, intraspecies sample comparison tool for sample swap detection. Gigascience 13, giae024 (2024).

7. Haas, D. M. et al. A description of the methods of the Nulliparous Pregnancy Outcomes Study: monitoring mothers-to-be (nuMoM2b). Am. J. Obstet. Gynecol. 212, 539.e1-539.e24 (2015).

8. Austin, G. I. et al. Contamination source modeling with SCRuB improves cancer phenotype prediction from microbiome data. Nat. Biotechnol. 41, 1820–1828 (2023).

9. Austin, G. I. & Korem, T. Planning and analyzing a low-biomass microbiome study: A data analysis perspective. J. Infect. Dis. jiae378 (2024).

10. Goltsman, D. S. A. et al. Metagenomic analysis with strain-level resolution reveals fine-scale variation in the human pregnancy microbiome. Genome Res. 28, 1467–1480 (2018).

11. Minich, J. J. et al. Quantifying and understanding well-to-well contamination in Microbiome research. mSystems 4, (2019).

12. Austin, G. I. et al. Processing-bias correction with DEBIAS-M improves cross-study generalization of microbiome-based prediction models. Nat. Microbiol. 1–15 (2025).

13. Kindschuh, W. F. et al. Early prediction of preeclampsia using the first trimester vaginal microbiome. bioRxiv 2024.12.01.626267 (2024) doi:10.1101/2024.12.01.626267.

14. Khan, R. R. et al. Genetic polymorphisms associated with adverse pregnancy outcomes in nulliparas. Sci. Rep. 14, 10514 (2024).

15. Bolger, A. M., Lohse, M. & Usadel, B. Trimmomatic: a flexible trimmer for Illumina sequence data. Bioinformatics 30, 2114–2120 (2014).

16. Langmead, B. & Salzberg, S. L. Fast gapped-read alignment with Bowtie 2. Nat. Methods 9, 357–359 (2012).

17. Li, H. et al. The Sequence Alignment/Map format and SAMtools. Bioinformatics 25, 2078–2079 (2009).

18. Perez, G. et al. The UCSC Genome Browser database: 2025 update. Nucleic Acids Res. 53, D1243–D1249 (2025).

